# Evidence of an optimal error rate for motor skill learning

**DOI:** 10.1101/2023.07.19.549705

**Authors:** Naser Al-Fawakhiri, Sarosh Kayani, Samuel D. McDougle

## Abstract

When acquiring a motor skill, learners must practice the skill at a difficulty that is challenging but still manageable to gradually improve their performance. In other words, during training, the learner must experience success as well as failure. Does there exist an optimal proportion of successes and failures to promote the fastest improvements in skill? Here, we build on a recent theoretical framework for optimal machine learning, extending it to the learning of motor skills. We then designed a custom task whose difficulty dynamically changed along with human subjects’ performance, constraining the error rate during training. In a large behavioral dataset (N=192), we observed evidence that learning is greatest at around a ∼30% error rate, matching our theoretical predictions.

## Introduction

Acquiring a new motor skill involves reducing errors (Krakauer & Mazzoni, 2011; Ohlsson, 1996; Tseng et al., 2007). Consider learning to shoot free throws in basketball: After a miss, the player uses the observed error information (e.g., “I missed to the right”) to fine-tune future attempts (e.g., re-orienting the shot slightly left). Such performance errors during motor skill learning drive improvements in both accuracy and precision and, given enough time and practice, can lead to automaticity and expertise (Haith et al., 2022; Shmuelof et al., 2012; Yang et al., 2022).

Importantly, one’s error rate during training can relate to if, and how well, they learn a skill. In the extreme, if one *never* experienced success, their motor system may have no template upon which to improve their skill (Sanger, 2004). On the other hand, learning would be unnecessary if one always experienced success (e.g., the hoop moved to wherever the ball was thrown). Thus, it is reasonable to assume that there is a “sweet spot” error rate that may generate the most efficient learning process (Guadagnoli & Lee, 2004). Indeed, in the context of simple non-motor skills (e.g., perceptual decision-making) it has recently been proposed that an optimal proportion of success and failure – an optimal “error rate” – can maximize the learning rate of a perceptual skill, such as binary categorization of sensory input (Wilson et al., 2019). Here, we extend this logic to the domain of sensorimotor skill learning.

In any task, errors become more likely as the difficulty of the task increases. Since the difficulty of the task can often be externally modulated, controlling an individual’s error rate during training requires dynamically controlling the difficulty of the task in response to their performance, exemplified by classic psychophysical methods like “staircasing” (Cornsweet, 1962). But how does task difficulty influence learning? Wilson and colleagues (2019) argue that learning using a gradient descent algorithm mathematically specifies an optimal error rate with very few additional assumptions. The authors considered the context of a binary discrimination task and, using their framework, predicted an optimal error rate of ∼15%, a value that was validated by multiple simulations over a range of binary decision-making tasks.

Can a similar framework be applied to motor skill learning? The first relevant issue is the putative relationship between motor skill learning and gradient descent. It has been suggested that motor skill learning, at least in the form of gradual improvements in skill that reduce errors and improve precision (Diedrichsen & Kornysheva, 2015; Sternad, 2018), does indeed operate via a gradient descent process (Greenstreet et al., 2025; Haith, 2025; Pierella et al., 2019). In many motor skills, the learner begins with an initially noisy policy and gradually refines this policy, reducing the variability of their actions as their skill improves (Haith, 2025; Sternad, 2018). For example, a novice darts player may make errors that span the whole width of the dartboard initially. But, as they improve, they gradually decrease their variability to hone in on a particular region of the board.

Fitts & Posner (1967) proposed a three-stage model of motor learning: the cognitive, associative, and autonomous stages. In the cognitive stage, learners must learn the goals of the task and the appropriate strategies to adopt to achieve those goals, heavily relying on explicit knowledge to achieve success in the task. In the associative stage, the learner begins to refine their actions, steadily reducing execution errors and variability. Finally, in the autonomous stage, the learner’s behavior has been practiced so thoroughly as to become automatic, requiring little cognitive effort.

To return to the darts analogy, an early novice may begin in the cognitive stage by trying to figure out the appropriate movements to launch a dart in the general direction of the dartboard. This process involves figuring out the appropriate strategy to launch the dart and aim at a particular spot on the board. After learning these strategies, the learner may eliminate any initial biases, but he will still have highly variable motor output around the region of the board he is aiming at. The subsequent associative stage marks a period of refinement, or precision improvement, where he steadily reduces that variability to hone in on the desired region of the board. It is on this stage of learning that this paper focuses.

Here, we hypothesized that the framework proposed by Wilson and colleagues (2019) might apply to motor skill learning of the form described above, even though motor errors are both signed and continuous (rather than binary and unsigned, as in perceptual discrimination tasks). First, we analytically extended their model to demonstrate that a roughly 32% error rate may be the optimal error rate for motor skill learning, given some basic assumptions. After specifying this prediction, we developed a customized “Pong” task that required precise online visuomotor control and collected behavioral data on the task from a large sample of human participants (N = 194). Using a custom dynamic error-control algorithm, we modified the size of subjects’ Pong paddle on a trial-by-trial basis to impose different error rates on their performance and measured their improvements in the task.

Strikingly, we found reliable evidence that individuals with an error rate around ∼32% showed the most learning in the task, directly in line with our model simulations. The current work provides a theoretically grounded specification of a putative optimal error rate for motor learning, and may thus inform both practical applications (e.g., sport pedagogy, clinical rehabilitation protocols) and theories concerning the underlying neural algorithms for motor skill learning.

## Methods

### An Optimal Error Rate Framework for Motor Skill Learning in One Dimension

In an idealized motor learning task, an agent prepares an action ***a*** in response to some set of internal conditions or policy ***π*** and external stimuli or state ***s***. Thus,

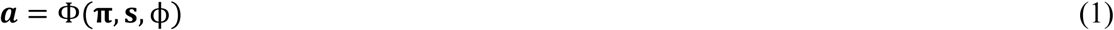

where Φ(⋅) represents the function that transforms ***π*** and ***s*** into motor actions, and ϕ is a set of internal, tunable free parameters trained by learning.

Whenever an action is executed, it is evaluated in terms of the discrepancy between the actual movement outcome a and the intended outcome i. This discrepancy is a motor error ***e***

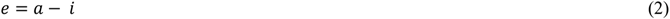

Here, motor learning consists of minimizing this error (distance from the center of a dart board, a golf hole, or a basketball hoop; in our Pong task, the distance of the puck from the center of the paddle). We further assume that errors are normally distributed around the target action and that there is no systematic bias (i.e., the mean of the policy is the optimal solution). In the Pong task, the trial ends when the participant intercepts the puck, and participants are instructed to adopt a strategy of aiming to intercept the puck directly in the middle of the paddle to afford themselves the highest chance of intercepting it (i.e., participants are explicitly provided the strategy that would result in errors with zero-mean). Thus, ***e*** can be rewritten as a Gaussian with zero-mean and a standard deviation of σ

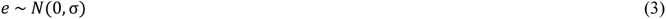

While we assume a normal distribution, any symmetrical, zero-mean error distribution can be used and will converge to an optimal error rate that depends on the specific choice of error distribution (see Wilson et al. 2019).

Both laboratory and naturalistic motor tasks typically have some degree of error tolerance, Δ, which still permits task success despite noise. This tolerance can take many forms: the radius of the green or the hole in golf, the radius of the dartboard when shooting darts, the width of the basketball hoop; or, in our Pong task, the center-to-edge length of the paddle.

Under these conditions, the error rate at any point in the task can be formalized as:

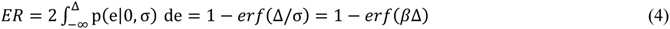

where *p*(⋅) is the standardized error distribution, here a normal distribution, *erf*(*x*) is the Gaussian error function, and β = 1/σ is the agent’s motor precision. In our case, β would be the inverse of the root-mean-squared-error (RMSE) centered around the intended outcome ***i*** (i.e., the center of the paddle in Pong, or the center of the target).

The goal of learning is to reduce error, which would require tuning the parameters ϕ in order to generate an action ***a*** as close to ***i*** as possible. This amounts to tuning the parameters of the policy to increase the precision of the policy around the optimal solution, thereby increasing β (Greenstreet et al., 2025; Haith, 2025; Sternad, 2018). Assuming tuning of these parameters is done through a gradient descent process, we can formalize the learning process over time as:

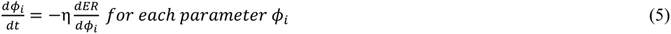

where η is the stepwise learning rate of the gradient descent algorithm. Given Equation 4, the derivative of the error rate 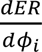 can be rewritten via the chain rule:

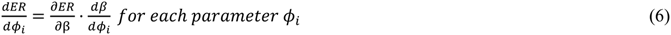

The first term of the right-hand side is the only term which depends on Δ, the task error tolerance (which can be experimentally controlled), as the second term only reflects how precision changes with the tuning of the relevant internal parameters outside of experimental control. Thus, to maximize the learning rate in Equation 5, we must maximize 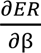 by optimizing the task error tolerance Δ at each time point *t* in training (see *Methods*). Using Equation 4, 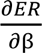 is given by:

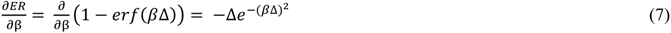

To maximize Equation 7, we must find its critical points by taking the derivative with respect to the error tolerance Δ and setting this derivative to zero:

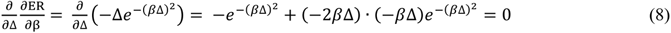

Simple algebraic manipulation of Equation 8 leads to:

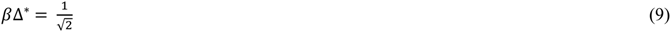

which can be plugged directly into Equation 4 to solve for the optimal error rate:

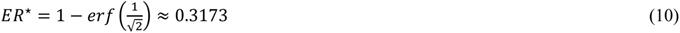

In other words, in this framework, the optimal error rate for motor learning in a one-dimensional task is 31.73%, and an optimal training schedule should aim to maintain performance at a 68.27% success rate throughout.

### Participants

200 participants (N=50/group; 46% female; age: 27±5) were recruited online through Prolific. Sample size was not determined by a-priori power analysis; instead, this sample size was chosen to ensure adequate inter-subject variation in error rates for our final analysis, which required subjects to complete the task with a wide range of experienced error rates. Recruitment was restricted to right-handed or ambidextrous individuals in the United States between the ages of 18 and 35, who had at least 40 prior Prolific submissions. All participants provided informed consent, in accordance with procedures approved by the Yale University Institutional Review Board (Protocol #2000027351 Psychological Mechanisms of Skill Learning). Due to participant data not posting to the database, data was not recorded from two participants. Four participants were further excluded for not making any recorded movements during the task. In analyses of learning indices (see *Statistical Analyses*), subjects whose learning index deviated by 3 times the group standard deviation from the group mean were excluded. This excluded 2 subjects. The final experimental sample was 192 subjects.

### Task Design and Protocol

Participants were recruited on Prolific to play a modified version of the game Pong, written using PhaserJS 3 (phaser.io). Participants controlled a 15-pixel wide rectangular paddle, initially set to 200 pixels (px) in length and fixed 10 px from the left edge of the workspace, inside a 1000 px by 800 px arena enclosed within a white border. Movements of subjects’ mouse or trackpad in the coronal plane directly translated to displacements of the paddle’s vertical position (i.e., movements away from the body = up; toward = down). Participants were instructed to move the paddle to intercept a small circular puck (14 px diameter) which would move toward their paddle at an angle, bouncing off the walls of the arena. They were specifically instructed to try to cause the puck to hit the middle of the paddle, as this is the strategy most likely to result in success. This task required participants to integrate visual sensory information regarding the position of the puck into their movements of the paddle in order to successfully intercept the puck.

Each trial consisted of the puck crossing from 25 pixels left of the right side of the arena to the left side of the arena where the participant’s paddle was located, crossing a horizontal distance of 975 px. At the start of each trial, the paddle’s vertical position was always centered, and the puck began at a random vertical position sampled from a uniform distribution extending ±375 px from the center of the screen. The launch angle of the puck could be ±30-45°, to ensure that the trajectory of the puck was unpredictable. The puck began moving 1000ms after the start of the trial. If participants successfully intercepted the puck, a green “+10” text prompt would appear in the center of the screen for 1000 ms, indicating success. Otherwise, if the puck hit the edge of the arena without being intercepted by the paddle, a red “+0” text prompt was displayed in the center of the screen for 1000 ms, indicating failure. A running points total was displayed above the arena and participants were told to maximize the points they earned. After presentation of reward feedback, the paddle was re-centered, and the puck appeared in a new location to start the next trial.

### Calibration and Error-control Algorithm

The experiment began with a 20-trial calibration phase, in which the horizontal speed of the puck was calibrated to be comfortable for each participant. The vertical speed of the puck was determined by the launch angle on a given trial. Since the duration of each trial was dependent on the horizontal speed of the puck, we needed to calibrate this speed to ensure every participant had ample time to react to the puck and adjust the paddle. The speed was calibrated using a staircase procedure: if the participant failed to intercept the puck, the horizontal speed was reduced by 75 px/s; if the participant successfully intercepted the puck, the speed increased by 75 px/s. The speed was initialized at 800px/s. The minimum allowed speed was 100px/s, but this was never achieved by any participant. This procedure resulted in a 72% (SD: 12%) success rate during the calibration phase.

Since this calibration phase often ended with a speed that slightly exceeded the comfortable range for the participant, the speed was cut to 80% of the final speed achieved. At this point, a 10-trial baseline began in which the puck speed and the paddle size were held constant. This allowed us to assess each participants’ initial degree of skill in the game. Mean success was 77% (SD:17%) and was not different between groups. At this point, the final 70-trial training phase began, and the paddle size was allowed to dynamically update according to a custom algorithm that monitored the recent distribution of errors to constrain participants’ error rates. Each participant was randomly assigned to one of four groups, each with their own intended error rate: 15%, 30%, 50%, and 65%. On every trial, the puck’s final vertical position relative to the final vertical position of the center of the paddle was added to a sliding 20-trial buffer, regardless of whether the trial was successful or not. After excluding outliers (defined as 3 times the median absolute deviation from the median; this is identical to the MATLAB *rmoutliers* function), we computed the root-mean-squared-error (RMSE) of this 20-trial history. From this we inferred a zero-mean Gaussian distribution with the RMSE as the standard deviation and, using the CDF of the Gaussian distribution, we computed a paddle size that would result in the intended error rate. This outlier criterion was chosen to only detect extreme outliers that could vastly impact the RMSE calculation for the duration of the 20-trial buffer, as determined by pilot experiments.

### Statistical Analysis

The primary dependent measure on each trial was the task error, defined as the difference between the final vertical position of the puck and the final vertical position of the center of the paddle. Errors from every ten trials were binned to compute the RMSE, and these RMSEs were inverted in order to estimate precision, β. We note that the overall pattern of results does not change if the inverse of the standard deviation of the errors, an alternative measure of precision, was used instead of the inverse RMSE. RMSE was preferred since participants were instructed to aim to intercept the puck in the middle of the paddle, so any biases were random and unintentional. Each subject’s baseline (trials 21-30) precision was subtracted from every subsequent bin in the training phase in order to quantify improvement. Average improvement, or the learning index (LI), was quantified as the average β of all training bins after baseline correction. Statistical outliers, defined as 1.75 times the interquartile range (IQR) away from the median of each of the four error rate groups, were excluded. This excluded 10 people (5.15%), leaving us with a final experimental sample of 184.

### Theoretical Modeling

The theoretical predictions plotted in Figure 1 and Figure 3D were computed in the manner described above. Since the integral of a Gaussian from −∞ to *x* cannot be analytically solved, it was estimated using a 100,000-rectangle Riemann sum from -10 to *x*, a computationally costly but highly accurate estimate of the integral. 30,000 values of Δ from 0 to 3 in increments of 0.0001 were used to generate the traces in Figure 1 as well as to generate a function that translated Δ values into theoretically predicted learning rates 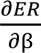, using the *approxfun* function in R (Becker, 2018). Since the units of these predicted learning rates are arbitrary, we fit scaling and offset free parameters that allowed the predictions to match the observed precision improvements of our subjects using R’s optim function most closely, and the resulting trace was plotted in Figure 3C.

**Figure 1.**
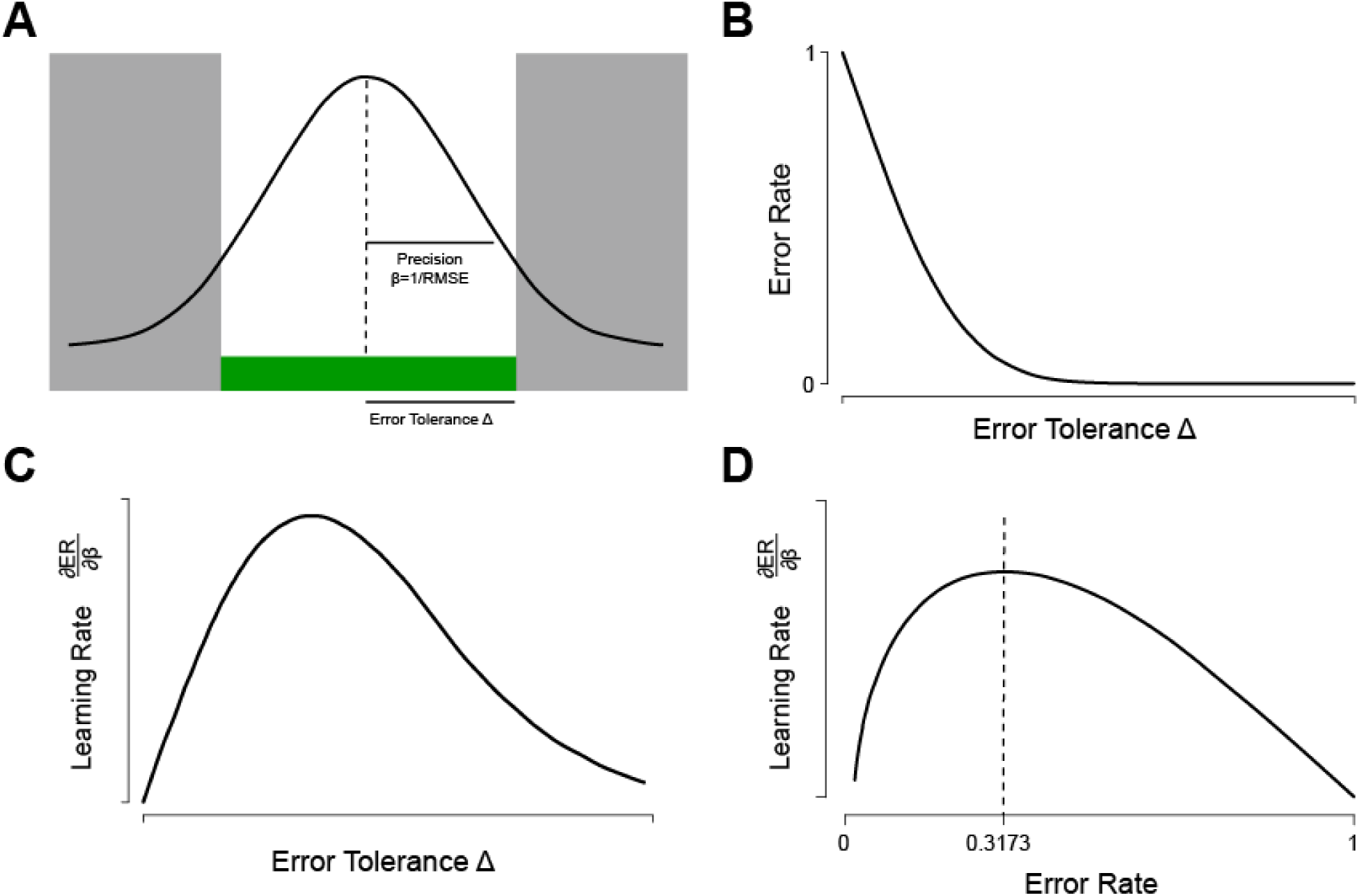
Derivation of an optimal error rate for motor learning. Figure adapted from Wilson et al. (2019). (A) Probability distributions of a given motor error *e*, within a certain error tolerance Δ. Shaded area represents the probability of a failure, or the error rate. Across training, the magnitude of errors shrinks, resulting in increased precision β. (B) As the error tolerance Δ increases (i.e. decreasing difficulty), fewer actions result in failures, leading to a decrease in the error rate. (C) The learning rate is given by the derivative of the error rate (ER) with respect to the agent’s precision β. Maximal learning occurs at an error tolerance Δ^∗^which changes with skill/training (β). (D) Same as C but replotted with the corresponding error rates on the x-axis. The optimal error rate remains constant, regardless of precision β, at ∼31.73%.

## Results

### Participant performance in one-dimensional Pong adhered to key theoretical assumptions

We aimed to test our theoretically grounded “optimal error rate” for motor learning using a motor skill task fashioned after the video game Pong, where participants (N=192) controlled the vertical position of a rectangular paddle using a mouse or touchpad to intercept a small white puck that moved across the screen at various angles.

Participants were instructed to aim for the middle of the paddle, as this would offer them the greatest chance of successfully intercepting the puck on each trial. This aligned with ensuring that participants adopted a strategy to minimize their errors from the center of the paddle, and, on average, had errors with zero mean bias. The mean error from the center of the paddle across all participants on all trials after the familiarization phase was 1.90 pixels (SD: 102 px) prior to removing outliers (>3SD from the mean), and 0.84 px (SD: 93 px) after removing outliers. This was not statistically significantly different from 0 (t(15185)=1.12, p=0.26). Analyzing each individual subject’s mean errors, participants averaged errors of 0.64 px from the center of the paddle (SD across participants: 12 px; outliers removed). When tested against zero using a Student’s t-test, only 7.2% of participants had errors that significantly differed from zero.

Another key assumption of our model was that errors were normally distributed. The overall distribution of errors across all trials and all participants, after removing outliers, exhibited a skewness of -0.077 and kurtosis of 0.70. When each participant’s distribution of errors was tested for normality using a Shapiro-Wilk test, only 11% of participants had errors that significantly deviated from the normal distribution. Together, 31 participants (16%) had either non-zero-mean errors or non-normally-distributed errors, and excluding these participants did not change the overall pattern of our results.

A final assumption was that learning proceeded via gradient descent; in other words, learning proceeded such that participants made actions on subsequent trials to reduce their errors by a proportion of the error observed on preceding trials (as is the case in classical state-space models in motor learning; state-space models are a variant of gradient descent). State-space learning models, and gradient descent learning rules in general, are well established in modeling motor learning in a variety of tasks (Thoroughman & Shadmehr 2000; Taylor & Ivry 2011; McDougle et al. 2015; Krakauer et al. 2019; Pierella et al. 2019). Here, we employed a linear mixed-effects model to fit each subject’s error magnitude across trials using a linear model, a quadratic model, and an exponential model. The exponential model is predicted by gradient descent if participants reduce their errors on subsequent trials by a proportion of their errors on preceding trials. Model comparison by BIC showed that the exponential model best fit the data (ΔBIC linear: -14.9, ΔBIC quadratic: -0.9), but the quadratic fit could not be rejected. Model performance by R^2^ favored the exponential fit (exponential: 0.10, quadratic: 0.04, linear 0.08). We note the low R^2^ values are due to fitting each individual trial’s error, which is remarkably noisy. Given the well-established nature of the use of gradient descent models in motor learning, we believe this data is consistent with the assumption that learning proceeded by gradient descent in our task.

### Skill improvement in a one-dimensional motor task is optimized when performance is maintained near a 30% error rate

To scale the difficulty of the task with participants’ evolving skill level and thus attempt to maintain a fixed error rate during training, the paddle’s size dynamically changed across trials according to the distribution of participants’ recent errors (see *Methods* for details). Participants were randomly sorted into 4 groups, each with an imposed error rate: 15%, 30%, 50%, and 65%. It should be noted that, while the dynamic paddle size aimed to constrain participants’ error rates to these pre-specified levels, variation in error rate was common, and each participant indeed experienced their own unique error rate during training (Fig. 2B). On average, across the 70-trial training phase, the dynamic algorithm undershot the intended error rate by 3.1% (SD: 6.6%), or 2.2 (SD: 4.6) trials. The 65% error rate group did have modestly, but significantly greater, success than the intended 35% success rate (t(48)=3.03, p=0.004, average: 41.6% success, SD: 6.1%). These slight deviations were ultimately inconsequential, as our true goal was to produce a wide range of error rates during training to correlate each participant’s learning rate with the learning rate theoretically predicted for their individual error rate. In this goal, our algorithm succeeded. Participants in the 15% error rate group had error rates ranging from 5% error to 31%. Participants in the 30% error rate group had error rates ranging from 14% to 40%. Participants in the 50% group had error rates that ranged from 28% to 55%. Participants in the 65% group had error rates that ranged from 38 to 65%.

**Figure 2.**
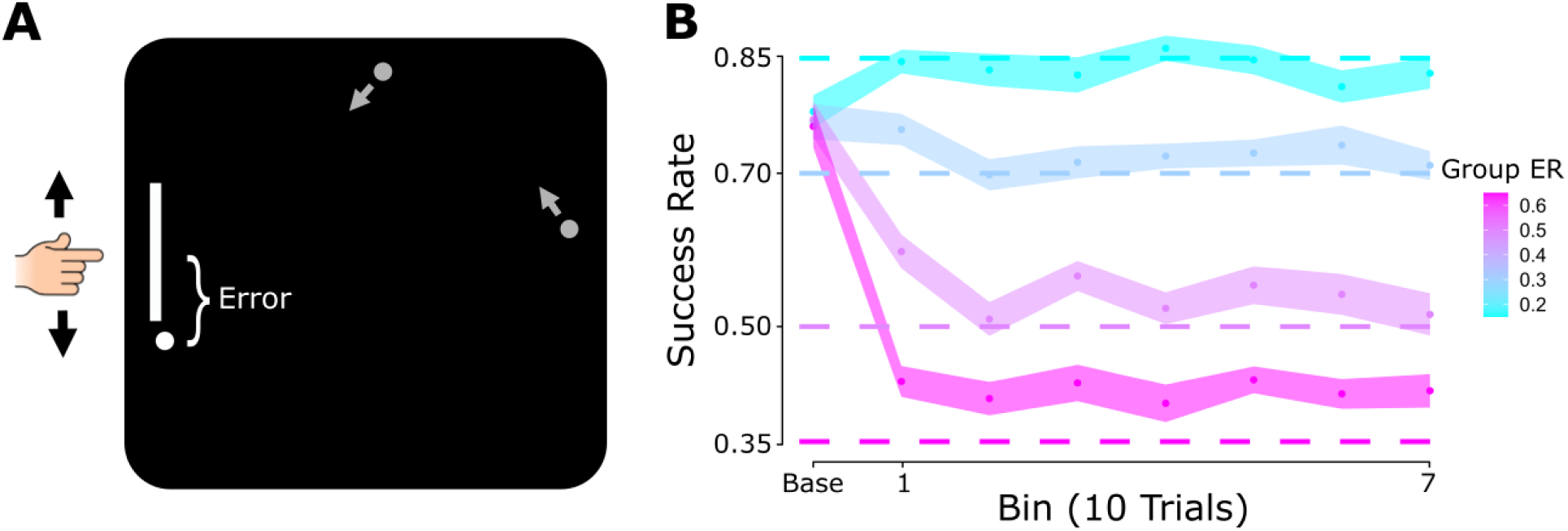
Task description. (A) Participants played a modified version of Pong, where they could control the paddle’s vertical position with their mouse in order to intercept the moving puck. (B) The paddle’s length was dynamically controlled on each trial by an algorithm designed to control the error rate across trials. Each participant was randomly assigned to one of four groups, each with a specific error rate (“ER”; 15%, 30%, 50%, 65%). The algorithm was largely successful in limiting the success rates of participants in each group. Shading reflects mean ± 1 S.E.M.

After an initial 20-trial calibration phase and 10 baseline trials (see *Methods*), participants completed 70 training trials in which the paddle size varied with their performance. To track performance improvements across the task, we computed the root-mean-squared-error (RMSE) of the puck’s final position relative to the center of the final position of the user-controlled paddle for every 10-trial bin. As a proxy for precision, we inverted this RMSE to yield β – β is low when precision is low, and high when precision is high. Since every participant entered the task at various degrees of skill, we corrected these β values using the 10-trial baseline in order to measure each participant’s *improvement* (i.e., learning) across the length of the task. A “learning index” (LI) was quantified as the average β over all 70 training trials.

Across all error rate groups, participants improved their precision (inverse RMSE, β) during the training phase (Fig. 3A). Participants’ precision in the final 10 trials was significantly greater than their precision in the first 10 training trials (t(193)=5.68, p=4.8×10^-8^, d=0.41), and participants’ overall improvement, or Learning Index (LI; β averaged across all 70 training trials) was significantly greater than zero (t(193)=4.33, p=2.4×10^-5^, d=0.31).

**Figure 3.**
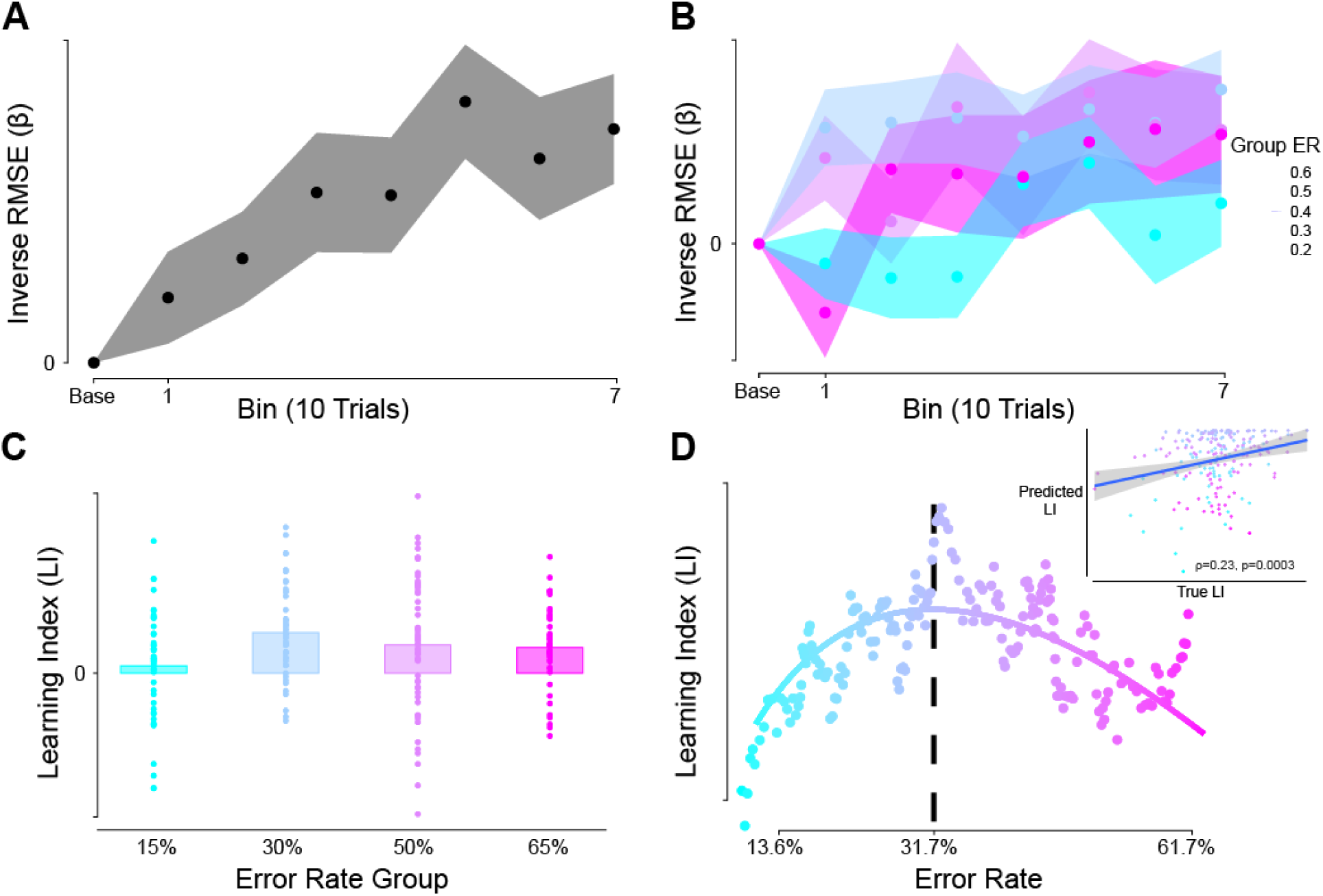
Optimal learning when training has an error rate of ∼31.7%. (A) All participants significantly improved over the course of training. β values are the baseline-corrected inverse RMSE for each 10-trial bin (arbitrary units). (B) Each group displayed different degrees and time courses of improvement. (C) Learning index reflects the average β across all 7 training bins for each subject. The group with the intended learning rate of 30% exhibited the greatest learning. (D) Individual subjects’ error rates plotted against their individual learning index. The smooth curve reflects the theoretically predicted learning rates for each error rate. *Inset:* Spearman correlation between subjects’ true LI and theoretically predicted LIs for their experienced error rate during training. Shading in panels A & B and error bars in panels C reflect standard errors of the mean (SEM).

Different error rate groups exhibited different time courses and degrees of improvement (Fig. 3B, C). As predicted, the 30% error rate group exhibited the greatest LI over all four groups: Their LIs were significantly greater than the 15% group (t(89)= 3.11, p=0.003, d=0.65), and marginally greater than the 65% group (t(87)=1.58, p=0.12, d=0.33), but not significantly greater than the 50% group (t(94)=0.98, p=0.32, d=0.20). Critically, as noted above, group-wise comparisons were not particularly appropriate given individual variation in actually realized error rates within each group.

Our main analysis involved comparing each participant’s LI to the theoretical learning rate predicted by our model (Fig. 3D). Supporting our theoretical predictions, participants’ learning rates significantly correlated with the learning rate predicted for their experienced error rate during training (Pearson correlation: R=0.25, p=0.001; Spearman correlation: *ρ*=0.23, p=0.003). This is a remarkable result given that our model does not require any fit parameters to achieve this degree of correlation. The correlation remained significant after removing participants with non-zero-mean errors or errors that were not normally distributed (Pearson correlation: R=0.26, p=0.001; Spearman correlation: *ρ*=0.23, p=0.004).

In conclusion, our theoretical prediction of an optimal motor skill learning error rate of ∼32% was supported in a large dataset consisting of people performing and improving at a continuous visuomotor skill task. These data are consistent with the theoretical framework proposed by Wilson et al. (2019), here generalized to the novel case of motor learning.

## Discussion

Here, we developed a new theoretical framework on optimal motor skill learning and employed a novel Pong-based motor skill task to provide behavioral evidence supporting our theoretical prediction that the optimal error rate for learning this and similar (one-dimensional) motor skills should be ∼32%. By controlling human subjects’ error rates during our Pong task, we could influence how well they learned throughout the course of training. Our theoretical predictions, which were closely based on recent work on optimal learning for binary classification tasks (Wilson et al., 2019), were supported both by showing the strongest learning in the group with the near-optimal error rate and by revealing a significant correlation between the model predictions and the empirical data across a range of individuals’ actual error rates (Figure 3). We propose that if an error rate of approximately ∼32% is maintained throughout training of a simple (one-dimensional) motor skill, one may expect to observe the greatest improvements in said motor skill. This discovery echoes work on “error-less” learning paradigms, where high success rates are maintained throughout training, and the difficulty of the task increases as the learner improves (Capio et al., 2013; Maxwell et al., 2001).

### Why does the error rate during training influence learning?

In the mathematical framework proposed by Wilson and colleagues (2019) and extended here, the relationship between training difficulty and learning is a necessary consequence of tuning the parameters responsible for generating responses that reduce errors via gradient descent. Such an approach leaves a key question unanswered: how does training difficulty result in changes in learning rate?

The “Challenge Point” framework (Guadagnoli & Lee, 2004) proposes that an agent’s performance at a motor skill is a source of information for future learning and action attempts. Highly expected outcomes, such as success when the task is easy, yield very little information as the agent can fully predict the consequences of their action. However, highly unexpected outcomes, such as success when the task is extremely difficult, are rare but can provide a high degree of information. The “optimal challenge point” occurs when the task difficulty is calibrated to ensure a moderately high probability of success and maximally informative feedback. The balance of maximizing both the expectation of success and the amount of interpretable information available for learning results in a fixed optimal error rate during training of some skills (Wilson et al., 2019).

Another psychological explanation focuses on the role of reward and motivation in error-based learning. Motivation to perform well in a task can drive increased sensitivity to errors (Bengtsson et al., 2009), which in turn can prompt faster learning (Wu et al., 2006). Additionally, rewarding feedback tends to increase motivation (Bissonette & Roesch, 2016) and can accelerate motor learning (Nikooyan & Ahmed, 2015). Thus, the recent history of successes and failures may prime an “optimal” degree of motivation, where errors occur in concert with rewarded trials, keeping the agent’s error sensitivity and motivation for success high. Put another way, if the task is too easy, learning would be minimal; but, if the task is too difficult, successes are too infrequent to generate the benefits to motivation and error sensitivity that are crucial for maintaining a high learning rate. The balance of these two opposing forces could also result in an optimal error rate for learning.

The mathematical framework employed here does not allow us to distinguish among these explanations, nor other higher-level psychological theories not mentioned here. The formation of internal models for motor learning requires obtaining information about how observed sensory outcomes relate to internally generated action plans (Abdelghani et al., 2008; Pierella et al., 2019; Thoroughman & Shadmehr, 2000) in line with the Challenge Point hypothesis (Guadagnoli & Lee, 2004). But reward feedback has also been shown to enhance motor learning (Galea et al., 2015; Izawa & Shadmehr, 2011; Nikooyan & Ahmed, 2015), supporting a motivational account as well. Thus, both explanations are plausible and may reflect how different brain regions (e.g., the cerebellum and basal ganglia, respectively) contribute to the fine-tuning of motor learning-related parameters (Greenstreet et al., 2025; Seidler et al., 2013). In general, we position our current findings at a more fundamental, algorithmic level.

### Extensions and limitations of the current framework

In our experiment, we chose a simple motor task where motor errors are generally assumed to be normally distributed. However, in some tasks, such as interval timing (Swanton & Matell, 2011), the distribution of errors may not be normal and may follow other known distributions. As noted by Wilson and colleagues (2019), an optimal error rate using non-Gaussian error distributions can be computed using the same equations described here, without guarantees that the optimal error rate derived in this manner does not depend on parameters of the error distribution. In tasks where the error distribution is unknown or cannot be mathematically described, empirical observation or alternative methods of estimating the distribution must be applied in order to compute the optimal error rate.

We also restricted our consideration to a simple one-dimensional motor skill task (errors were only in the vertical dimension). However, many motor tasks involve errors in multiple dimensions (e.g., a dart or archer’s arrow may miss the bullseye in both the vertical and horizontal dimensions), which must be concurrently trained. Future work must be done to extend the optimal error rate framework to these more complex, naturalistic tasks. An important question which must be considered is whether learning in each dimension is fully independent, or if learning in one dimension affects learning in the others; the answer to this question would allow us to better model the error reduction processes involved in learning complex motor skills when multiple dimensions must be concurrently controlled (Haith et al., 2022; Pierella et al., 2019). Additionally, distributions of errors in each dimension may not follow similar mathematical descriptions, requiring more complicated multi-dimensional integrals to optimize the error rate. Nonetheless, the general mathematical framework demonstrated here can, in theory, be applied to such multi-dimensional tasks.

Lastly, our task only examines motor learning as defined as the gradual reduction in errors and improvement of precision over time through practice, which arguably represents only one component of skill learning (i.e., improvements in “motor acuity”; Krakauer et al., 2019). Other components of motor learning, including motor adaptation, may be better modeled as changes in the mean of the policy, rather than gradual refinement of the precision of the policy (Greenstreet et al. 2025). Explicit strategies, especially early in learning, can cause a learner to abruptly reduce their errors or discover new solutions (Mazzoni & Krakauer, 2006; Taylor & Ivry, 2011), abruptly shifting the mean of the policy instead of gradually learning via gradient descent. Explicit strategies, short-term memory, and cognitive insights during skill learning likely arise from separate cognitive processes not considered here, though they still make important contributions to skill learning (Haith & Krakauer, 2013; Hillman et al., 2025; McDougle et al., 2016, 2022; McDougle & Hillman, 2025). The model presented here is primarily concerned with how the learner learns to decrease the standard deviation of the policy around this mean, assuming the mean has already been discovered, thereby improving precision and reducing task-relevant variability (Haith, 2025; Sternad, 2018).

### Future Applications

Our results demonstrate that the error rate during training affects the rate of improvement of a simple visuomotor skill. We think this insight might be broadly applicable across applied domains. Rehabilitation paradigms, for example, have already recognized the need to adjust the difficulty of practice to optimally challenge patients and maximize the therapeutic benefit of training sessions (Lee et al., 2023; Metzger et al., 2014). In sports coaching, the principle of progressively increasing the difficulty of practice has been widely applied and shown to facilitate improvements in skill (Marraffino et al., 2021; Wickens et al., 2013; Zahran et al., 2020). Optimistically, these approaches, in both sports pedagogy and rehabilitation, may benefit from the theory used here to mathematically uncover the sweet spot of training difficulty for maximum skill improvement.

### Summary

We demonstrate that an optimal error rate for performance improvements in a motor skill may exist and suggest that this optimal error rate is around ∼32%. By testing a large cohort of human participants in a novel motor skill task (a modified version of Pong), we also provide empirical support for this prediction and demonstrate the potential importance of maintaining a particular error rate throughout training to maximize performance improvements. This approach may have broader implications in other domains concerned with optimizing the acquisition, refinement, and rehabilitation of motor skills.

## Declarations

### Ethics Approval

This research was approved by the Yale University Institutional Review Board (Protocol #2000027351 Psychological Mechanisms of Skill Learning).

### Consent for Participation and Publication

All participants provided written informed consent for their participation and the publication of data produced from their participation.

### Availability of Data, Materials, and Code

This study was not pre-registered. The Pong task used in this experiment can be found at pong-baseball-fast.netlify.app. Error rate can be specified using the URL parameter “GROUP=XX”, where XX is the intended error rate as a percentage (default value is 30). Data and analysis code can be found at https://osf.io/ew726/overview?view_only=61050a15edb746d1998444bb903e0142.

## References

Abdelghani, M. N., Lillicrap, T. P., & Tweed, D. B. (2008). Sensitivity Derivatives for Flexible Sensorimotor Learning. Neural Computation, 20(8), 2085–2111. 10.1162/neco.2008.04-07-507

Becker, R. (2018). The New S Language. Chapman and Hall/CRC. 10.1201/9781351074988

Bengtsson, S. L., Lau, H. C., & Passingham, R. E. (2009). Motivation to do Well Enhances Responses to Errors and Self-Monitoring. Cerebral Cortex, 19(4), 797–804. 10.1093/cercor/bhn127

Bissonette, G. B., & Roesch, M. R. (2016). Neurophysiology of Reward-Guided Behavior: Correlates Related to Predictions, Value, Motivation, Errors, Attention, and Action. In E. H. Simpson & P. D. Balsam (Eds.), Behavioral Neuroscience of Motivation (pp. 199–230). Springer International Publishing. 10.1007/7854_2015_382

Capio, C. M., Poolton, J. M., Sit, C. H. P., Holmstrom, M., & Masters, R. S. W. (2013). Reducing errors benefits the field-based learning of a fundamental movement skill in children. Scandinavian Journal of Medicine & Science in Sports, 23(2), 181–188. 10.1111/j.1600-0838.2011.01368.x

Cornsweet, T. N. (1962). The Staircase-Method in Psychophysics. The American Journal of Psychology, 75(3), 485–491. 10.2307/1419876

Diedrichsen, J., & Kornysheva, K. (2015). Motor skill learning between selection and execution. Trends in Cognitive Sciences, 19(4), 227–233. 10.1016/j.tics.2015.02.003

Galea, J. M., Mallia, E., Rothwell, J., & Diedrichsen, J. (2015). The dissociable effects of punishment and reward on motor learning. Nature Neuroscience, 18(4), Article 4. 10.1038/nn.3956

Greenstreet, F., Geerts, J. P., Gallego, J. A., & Clopath, C. (2025). Why motor learning involves multiple systems: An algorithmic perspective (p. 2025.12.19.695526). bioRxiv. 10.64898/2025.12.19.695526

Guadagnoli, M., & Lee, T. (2004). Challenge Point: A Framework for Conceptualizing the Effects of Various Practice Conditions in Motor Learning. Journal of Motor Behavior, 36, 212–224. 10.3200/JMBR.36.2.212-224

Haith, A. M. (2025). Policy-Gradient Reinforcement Learning as a General Theory of Practice-Based Motor Skill Learning (p. 2025.10.17.682587). bioRxiv. 10.1101/2025.10.17.682587

Haith, A. M., & Krakauer, J. W. (2013). Model-Based and Model-Free Mechanisms of Human Motor Learning. In M. J. Richardson, M. A. Riley, & K. Shockley (Eds.), Progress in Motor Control (pp. 1–21). Springer. 10.1007/978-1-4614-5465-6_1

Haith, A. M., Yang, C. S., Pakpoor, J., & Kita, K. (2022). De novo motor learning of a bimanual control task over multiple days of practice. Journal of Neurophysiology, 128(4), 982–993. 10.1152/jn.00474.2021

Hillman, H., McClure, T. N., & McDougle, S. D. (2025). Linking motor working memory to explicit and implicit motor learning. Journal of Neurophysiology, 134(6), 2036–2046. 10.1152/jn.00105.2025

Izawa, J., & Shadmehr, R. (2011). Learning from Sensory and Reward Prediction Errors during Motor Adaptation. PLOS Computational Biology, 7(3), e1002012. 10.1371/journal.pcbi.1002012

Krakauer, J., Hadjiosif, A., Xu, J., Wong, A., & Haith, A. (2019). Motor Learning. In Comprehensive Physiology (Vol. 9, pp. 613–663). 10.1002/cphy.c170043

Krakauer, J. W., & Mazzoni, P. (2011). Human sensorimotor learning: Adaptation, skill, and beyond. *Current Opinion in Neurobiology*, Sensory and Motor Systems, 21(4), 636–644. 10.1016/j.conb.2011.06.012

Lee, H., Choi, Y., Eizad, A., Song, W.-K., Kim, K.-J., & Yoon, J. (2023). A Machine Learning-Based Initial Difficulty Level Adjustment Method for Balance Exercise on a Trunk Rehabilitation Robot. IEEE Transactions on Neural Systems and Rehabilitation Engineering, 31, 1857–1866. IEEE Transactions on Neural Systems and Rehabilitation Engineering. 10.1109/TNSRE.2023.3260815

Marraffino, M. D., Schroeder, B. L., Fraulini, N. W., Van Buskirk, W. L., & Johnson, C. I. (2021). Adapting training in real time: An empirical test of adaptive difficulty schedules. Military Psychology, 33(3), 136–151. 10.1080/08995605.2021.1897451

Maxwell, J. P., Masters, R. S. W., Kerr, E., & Weedon, E. (2001). The implicit benefit of learning without errors. The Quarterly Journal of Experimental Psychology, 54A(4), 1049–1068. 10.1080/713756014

Mazzoni, P., & Krakauer, J. W. (2006). An Implicit Plan Overrides an Explicit Strategy during Visuomotor Adaptation. The Journal of Neuroscience, 26(14), 3642–3645. 10.1523/JNEUROSCI.5317-05.2006

McDougle, S. D., & Hillman, H. (2025). Motor working memory. Trends in Cognitive Sciences, S1364-6613(25)00236-0. 10.1016/j.tics.2025.08.011

McDougle, S. D., Ivry, R. B., & Taylor, J. A. (2016). Taking Aim at the Cognitive Side of Learning in Sensorimotor Adaptation Tasks. Trends in Cognitive Sciences, 20(7), 535–544. 10.1016/j.tics.2016.05.002

McDougle, S. D., Wilterson, S. A., Turk-Browne, N. B., & Taylor, J. A. (2022). Revisiting the Role of the Medial Temporal Lobe in Motor Learning. Journal of Cognitive Neuroscience, 34(3), 532–549. 10.1162/jocn_a_01809

Metzger, J.-C., Lambercy, O., Califfi, A., Dinacci, D., Petrillo, C., Rossi, P., Conti, F. M., & Gassert, R. (2014). Assessment-driven selection and adaptation of exercise difficulty in robot-assisted therapy: A pilot study with a hand rehabilitation robot. Journal of NeuroEngineering and Rehabilitation, 11(1), 154. 10.1186/1743-0003-11-154

Nikooyan, A. A., & Ahmed, A. A. (2015). Reward feedback accelerates motor learning. Journal of Neurophysiology, 113(2), 633–646. 10.1152/jn.00032.2014

Ohlsson, S. (1996). Learning From Performance Errors. Psychological Review, 103(2), 241–262.

Pierella, C., Casadio, M., Mussa-Ivaldi, F. A., & Solla, S. A. (2019). The dynamics of motor learning through the formation of internal models. PLOS Computational Biology, 15(12), e1007118. 10.1371/journal.pcbi.1007118

Sanger, T. D. (2004). Failure of Motor Learning for Large Initial Errors. Neural Computation, 16(9), 1873–1886. 10.1162/0899766041336431

Seidler, R. D., Kwak, Y., Fling, B. W., & Bernard, J. A. (2013). Neurocognitive Mechanisms of Error-Based Motor Learning. In M. J. Richardson, M. A. Riley, & K. Shockley (Eds.), Progress in Motor Control (pp. 39–60). Springer. 10.1007/978-1-4614-5465-6_3

Shmuelof, L., Krakauer, J. W., & Mazzoni, P. (2012). How is a motor skill learned? Change and invariance at the levels of task success and trajectory control. Journal of Neurophysiology, 108(2), 578–594. 10.1152/jn.00856.2011

Sternad, D. (2018). It’s not (only) the mean that matters: Variability, noise and exploration in skill learning. Current Opinion in Behavioral Sciences, Habits and Skills, 20, 183–195. 10.1016/j.cobeha.2018.01.004

Swanton, D. N., & Matell, M. S. (2011). Stimulus compounding in interval timing: The modality–duration relationship of the anchor durations results in qualitatively different response patterns to the compound cue. Journal of Experimental Psychology: Animal Behavior Processes, 37(1), 94–107. 10.1037/a0020200

Taylor, J. A., & Ivry, R. B. (2011). Flexible Cognitive Strategies during Motor Learning. PLOS Computational Biology, 7(3), e1001096. 10.1371/journal.pcbi.1001096

Thoroughman, K. A., & Shadmehr, R. (2000). Learning of Action Through Adaptive Combination of Motor Primitives. Nature, 407(6805), 742–747. 10.1038/35037588

Tseng, Y., Diedrichsen, J., Krakauer, J. W., Shadmehr, R., & Bastian, A. J. (2007). Sensory Prediction Errors Drive Cerebellum-Dependent Adaptation of Reaching. Journal of Neurophysiology, 98(1), 54–62. 10.1152/jn.00266.2007

Wickens, C. D., Hutchins, S., Carolan, T., & Cumming, J. (2013). Effectiveness of part-task training and increasing-difficulty training strategies: A meta-analysis approach. Human Factors, 55(2), 461–470.

Wilson, R. C., Shenhav, A., Straccia, M., & Cohen, J. D. (2019). The Eighty Five Percent Rule for optimal learning. Nature Communications, 10(1), Article 1. 10.1038/s41467-019-12552-4

Wu, S.-W., Trommershäuser, J., Maloney, L. T., & Landy, M. S. (2006). Limits to human movement planning in tasks with asymmetric gain landscapes. Journal of Vision, 6(1), 5. 10.1167/6.1.5

Yang, C. S., Cowan, N. J., & Haith, A. M. (2022). Control becomes habitual early on when learning a novel motor skill. Journal of Neurophysiology, 128(5), 1278–1291. 10.1152/jn.00273.2022

Zahran, L., El-Beltagy, M., & Saleh, M. (2020). A Conceptual Framework for the Generation of Adaptive Training Plans in Sports Coaching. In A. E. Hassanien, K. Shaalan, & M. F. Tolba (Eds.), Proceedings of the International Conference on Advanced Intelligent Systems and Informatics 2019 (pp. 673–684). Springer International Publishing. 10.1007/978-3-030-31129-2_62

